# Genomics of experimental adaptive radiation in the cryptic coloration of feather lice

**DOI:** 10.1101/2024.12.20.629508

**Authors:** James Baldwin-Brown, Scott M. Villa, Emiko Waight, Kevin P. Johnson, Sarah E. Bush, Dale H. Clayton, Michael D. Shapiro

## Abstract

A major challenge faced by living organisms is adaptation to novel environments. This process is poorly understood because monitoring genetic changes in natural populations is difficult. One way to simplify the task is to focus on organisms that can be studied in captivity under conditions that remain largely natural. Feather lice (Insecta, Phthiraptera, Ischnocera) are host-specific parasites of birds that live, feed, and breed solely on feathers. Birds defend themselves against lice, which damage feathers, by killing them with their beaks during bouts of preening. In response, feather lice have evolved background-matching cryptic coloration to help them avoid preening. We experimentally manipulated the color backgrounds of host-specific pigeon lice (*Columbicola columbae*) by confining them to different colored breeds of rock pigeon (*Columba livia*) over a period of four years (ca. 60 louse generations). Over the course of the experiment, we sampled lice from pigeons every six months for genomic resequencing, and then calculated allele frequency differences and trajectories to identify putative genomic sites under selection. We documented many loci that changed in response to selection for color. Most loci putatively under selection were unshared among replicate populations of lice, indicating that independent adaptation of distinct lineages to the same novel environment resulted in similar phenotypes driven by different genotypes.

## 2 Introduction

Adaptive radiation occurs when members of a single lineage evolve different adaptive forms in response to divergent natural selection. Despite its fundamental importance to biodiversity, most of what we know about adaptive radiation is from historical inferences. Iconic examples include organisms that live on islands, such as Darwin’s finches, Caribbean *Anolis* lizards, and Hawaiian silversword plants (Schluter 2000). Islands are younger than mainland ecosystems, making it easier to reconstruct the history of colonization and speciation events for island radiations.

Some host-parasite interactions also provide tractable systems for the study of adaptive radiation because hosts are effectively living “islands” for host-specific parasites (Kuris et al. 1980). Parasitic feather lice feed on the downy regions of bird feathers, which reduces host survival and mating success (Clayton et al. 2016). Birds combat feather lice by removing them with their beaks during regular bouts of preening. Lice, in turn, have adaptations to avoid preening such as background-matching cryptic coloration: light colored species of birds host lighter species of lice, while dark colored species of birds host darker species of lice (Fig. 1; Bush et al. 2010).

**Figure 1:**
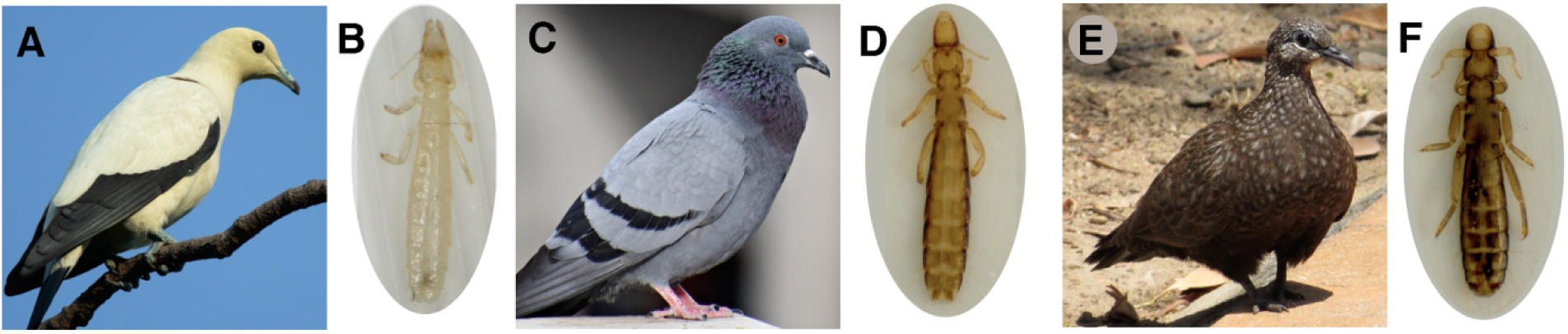
Examples of cryptic coloration among feather lice in the genus *Columbicola*. The Australian pied imperial-pigeon, *Ducula bicolor* (A), is parasitized by *C. wolffhuegeli* (B); the rock pigeon, *Columba livia* (C), is parasitized by *C. columbae* (D); and the chestnut-quilled rock pigeon, *Petrophassa rufipennis* (E), is parasitized by *C. masoni* (F). Photos by: (A) JJ Harrison, Wikimedia Commons, CC BY-SA 4.0; (B) SEB; (C) pxfuel.com/en/free-photo-qffys; (D and F) SMV and J. Altuna; (E) Nimzee, https://inaturalist.ala.org.au/photos/123753242.

Feather lice are permanent parasites that complete their entire life cycle on feathers (Clayton et al. 2016). A previous study (Bush et al. 2019) experimentally triggered diversification in the coloration of host-specific parasites that were confined to different colored host “islands.” The study employed an unusually tractable host-parasite system consisting of rock pigeons (*Columba livia*) and their feather lice (*Columbicola columbae*). Because the pigeon represents the entire habitat for its feather lice, populations of lice on captive birds live under conditions that are essentially natural. This fact, coupled with their short generation time (about 3.5 weeks from egg to adult, Martin 1934), makes feather lice good candidates for experimental evolutionary studies of adaptive radiation. Feather lice experimentally transferred to different colored rock pigeons experienced rapid and sustained divergence with populations of lice on white pigeons becoming lighter and populations of lice on black pigeons becoming darker over the course of the 48-month study (Fig. 2). Thus, the study identified the pattern and tempo of change in color among louse populations subjected to different host-imposed selective regimes.

**Figure 2:**
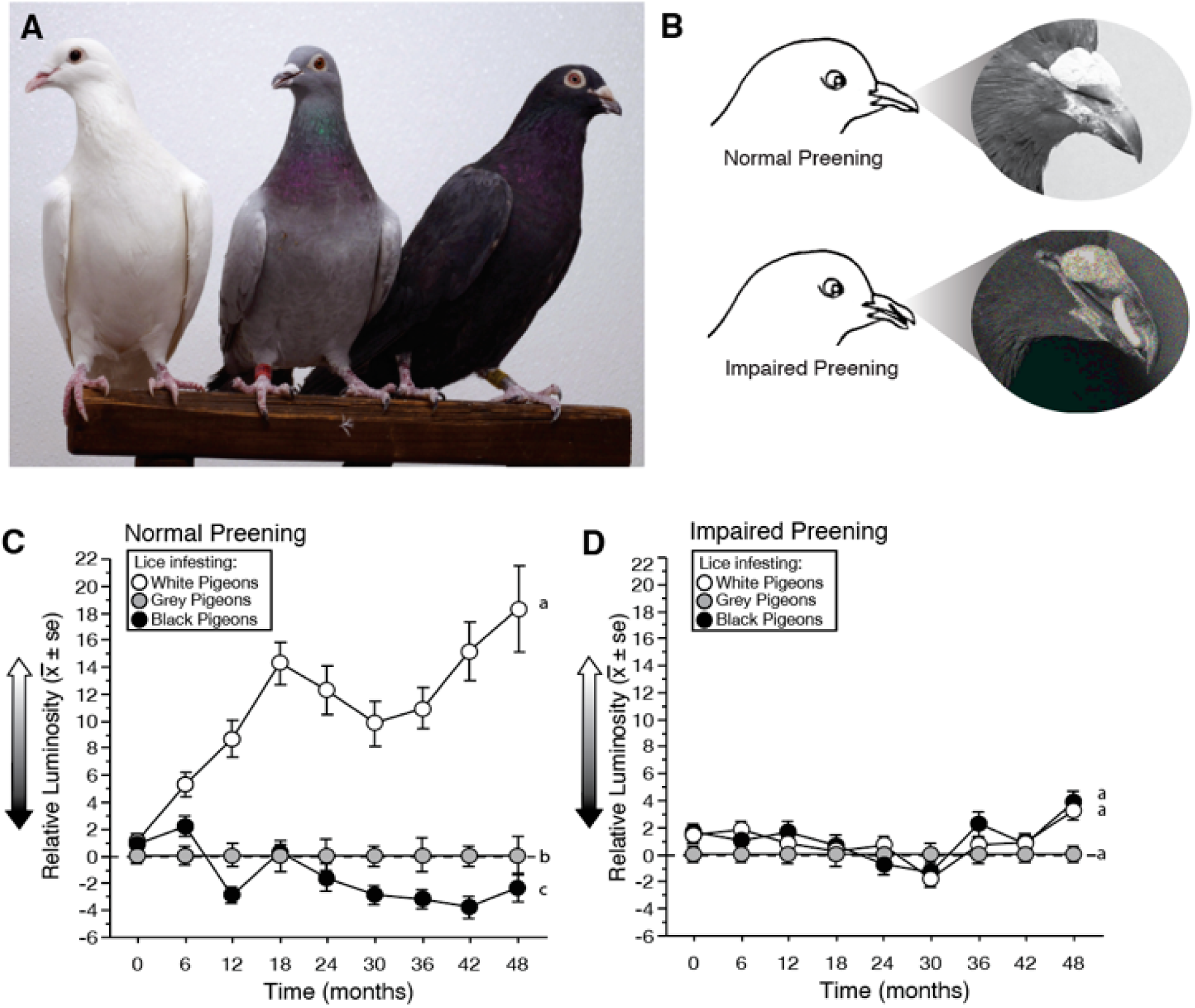
Experimental evolution of lice on hosts of varying color shows adaptation toward the host’s color. (A) Feather lice (*Columbicola columbae*) were transferred to three different rock pigeon color morphs: white (left), black (right), and grey (wild-type controls, middle). (B) Half of the pigeons could preen normally, while the other half were fitted with poultry “bits” to impair their preening ability. (C-D) Evolution of feather lice on different colored rock pigeons over a four-year period (ca. 60 louse generations). The y-axis shows changes in the mean (*±* se) luminosity (brightness) of lice on white and black rock pigeons, relative to lice on grey rock pigeon controls (set to zero). Different lower-case letters indicate statistically significant differences. (C) On birds with normal preening, the relative luminosity of lice on white pigeons increased rapidly (LMM, P < 0.0001); the relative luminosity of lice on black pigeons decreased, but more slowly (LMM, P = 0.001). (D) Relative luminosity did not change significantly over time on white or black pigeons with impaired preening (LMM, P *≥* 0.34 in both cases). (Panels B-D redrawn from Bush et al. 2019).

A key goal in studies of adaptative evolution is to understand the number and location of genetic factors contributing to changes in adaptive phenotypic traits, such as color. Therefore, in addition to quantifying phenotypic change over time, we sampled lice from each of four replicate lineages for each color selection regime for genomic analyses. Lice vary in color from light to dark, consistent with differences in melanin content observed in other insects. The melanin production pathway is well described in insects, especially in *Drosophila* spp. (True 2003). Thus, the many studies of melanin color and patterning diversity in insects provide numerous *a priori* candidate genes for adaptive color shifts in lice. Changes in some of these genes can produce large color shifts alone or in combination with a modest number of other genes (Zhang et al. 2017). Here, we use an “evolve and resequence” approach in which we sequenced samples of lice from selected and non-selected lineages of lice over the course of a 48-month experiment. The evolve and resequence approach has been used to identify loci underlying diversity in viruses, microbes, and metazoans (e.g. Wichman et al. 2000; Van Laere et al. 2003; Andersson and Hughes 2009; Barrick et al. 2009; Burke et al. 2010; Paterson et al. 2010; Rubin et al. 2010; Johansson et al. 2010; Turner et al. 2011).

The goal of the current study is to identify the tempo and mode of genomic change in populations of lice evolving cryptic coloration. Lice vary in color from light to dark, consistent with differences in melanin content observed in other insects. We investigate patterns of genome wide allele frequency changes in color selection lines of pigeon lice and evaluate whether putative targets of selection occur in regions that harbor melanin pathway genes. Prior studies provide some evidence that, when adaptation to a new optimal body color is repeated, the same genes are involved each time (Bilandžija et al. 2012; San Jose and Roulin 2018). In the current study we test whether the same or different genomic regions are involved in replicate color selection lines and find that largely different sets of loci change in parallel experimental manipulations.

## 3 Data Availability

Sequence data are available through NCBI SRA (BioProject PRJNA1038774). All analysis scripts are available through GitHub at https://github.com/jgbaldwinbrown/jgbutils.

## 4 Materials and Methods

### 4.1 Animal housing and husbandry design

We used lice from the experimental evolution study of Bush et al. (2019), the design of which is briefly summarized here. Prior to the start of the study, any resident “background” lice were eradicated by housing 96 captive pigeons under low humidity conditions (< 25% relative ambient humidity) for *≥* 10 weeks. This method kills lice and their eggs, while avoiding residues from insecticides (Harbison et al. 2008). Next, hundreds of lice were collected from wild caught feral pigeons. Random subsets of 25 lice were transferred to each of 96 captive pigeons housed in 24 aviaries (4 birds per aviary; aviary dimensions: 1.8 m Œ 1.5 m Œ 1.0 m). When lice were transferred to birds in the experiment, the relative humidity was increased to 35 - 60%, providing sufficient humidity for feather lice to survive and reproduce on the birds (Nelson and Murray 1971).

The captive pigeons consisted of 32 white birds, 32 black birds, and 32 “grey” birds (ancestral “blue-bar” phenotype, controls) (Fig. 2A). Preening, which is a pigeon’s major defense against ectoparasites (Clayton et al. 2005), was impaired using harmless poultry “bits”, which are C-shaped pieces of plastic inserted between the upper and lower mandibles of a bird’s beak (Fig. 2B). Bits have no apparent side effects and they do not impair the ability of birds to feed (Clayton and Tompkins 1995). Preening was impaired in half of the birds, chosen at random, with all birds in the same aviary assigned to the same preening treatment.

Each aviary housed two male and two female birds of a given color and preening treatment (Table 1). The inclusion of four birds per aviary kept birds socially active and helped to ensure that birds did not preen their lice to extinction, which can occur on isolated birds that double their preening rates out of apparent boredom (Waller et al. 2024). The *Columbicola columbae* lice used in our experiment are so specialized for life on feathers that they seldom, if ever, venture onto the host’s skin, or away from the host’s body. *C. columbae* requires direct contact between feathers to move between birds (Clayton and Tompkins 1994). Since contact was common among birds in each aviary, the lice in each aviary were considered members of a single population. Lice were not able to move between birds in different aviaries. *C. columbae* are also known to move between birds by phoretic hitchhiking on hippoboscid flies (Harbison et al. 2008); however, such flies were not present on any of the birds in our experiment.

**Table 1:**
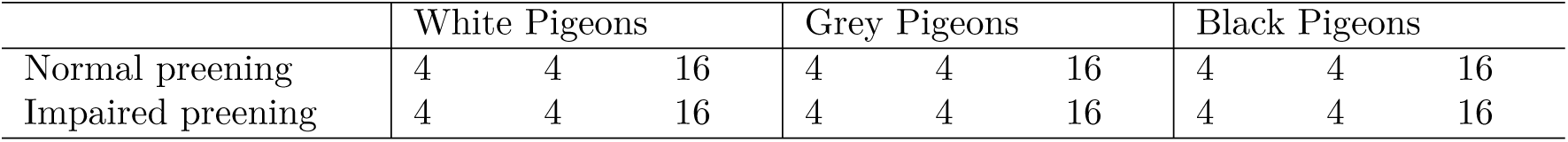
Distribution of birds among 24 aviaries in the experimental evolution study (Bush et al. 2019). Each aviary held two male and two female pigeons of the indicated color and preening treatment. Thus, there are four replicates in this experiment; each replicate consists of six populations of lice that include one population of lice from each color and preening treatment.

In summary, this experimental design included four replicates, each with six aviaries containing infested hosts of three colors and two preening treatments (Table 1). The 24 birds in each replicate were randomly seeded with lice from an infested captive population at the same time (as described above); however, the start dates of the six different replicates were staggered by approximately 6 weeks to allow time for the time-consuming activities associated with phenotypic and genomic sampling, described next.

### 4.2 Phenotyping and genomic sampling

At six-month intervals throughout the 48-month experiment, we removed random samples of lice from each population (Bush et al. 2019). Lice were removed by exposing birds’ feathers to CO2, which anesthetizes the lice. After exposure to CO2 the feathers of each bird were ruffled over a gridded sheet of paper on a tray to collect the anesthetized lice. The total number of lice removed from the four birds in an aviary was used to estimate louse population size. Up to 50 lice per population were chosen randomly from the grid, digitally photographed for phenotyping, then preserved at −80°C for genomic DNA extraction and sequencing. The remaining lice from the population were placed back on the birds. When fewer than 100 lice were recovered across the four birds in an aviary, only half of the louse population was chosen (randomly) for photographing and preservation. Thus, the number of preserved lice from each population ranged between a low of 5 and a high of 50.

For downstream analyses, we calculated the harmonic mean of census count for all time points for a population as a proxy for the effective population size (Supplementary Fig. 1). We used the harmonic mean to approximate the effective population size because it accounts for the effect of bottlenecks in the population size over time (Karlin et al. 1968).

### 4.3 Isolation of DNA

For collection timepoints at months 6, 12, 18, 24, 30, 42, and 48, we pooled samples from each louse population for DNA extraction, library preparation, and sequencing (3 colors, 2 preening conditions, and 4 replicates at 6 timepoints for a total of 144 sample pools). We also sampled and pooled 50 lice from infested wild caught feral pigeons each time a replicate was started (i.e., a time-matched sample of the ancestral population for each of the 4 replicates). Thus, we had a grand total of 148 pooled samples. We chose one time point, 36 months, for more intensive genomic analyses. This time point was chosen for two reasons: (1) phenotypic divergence had clearly occurred by that time, and (2) one of the populations (replicate 4 on white birds) began to decline in size at this time point. We sequenced lice individually at this time point to have the option to test for individual haplotypes that might be associated with regions under selection.

DNA from both pooled and individual samples was isolated by grinding with the TissueLyser LT (Qiagen) followed by DNA extraction with the Qiagen DNEasy Extraction Kit (Qiagen). Several modifications were made to the manufacturer’s protocol to extract DNA most effectively from the tough, chitinous lice. First, we ground the lice using seven one-minute TissueLyser cycles at the maximum frequency, then visually inspected the samples to confirm breakage of the cuticle. Second, we incubated tissue in lysis solution for between six hours and overnight. Third, for individually sequenced lice, we eluted DNA in 50 µL of elution buffer EB warmed to 65°C, then eluted again in a second 50-uL aliquot to maximize yield.

### 4.4 Genomic DNA sequencing

Libraries were constructed using high molecular weight DNA (1-20 ng) with the Nextera DNA Flex Library Prep kit (Illumina, Inc., San Diego, cat#20025520) and an average insert size of 450 bp. PCR-amplified libraries were quantified on an Agilent Technologies 2200 TapeStation using a D1000 ScreenTape assay (cat# 5067-5582 and 5067-5583); the molarity of adapter-modified molecules was defined by quantitative PCR using the Kapa Biosystems Kapa Library Quant Kit (cat#KK4824). Libraries were normalized and pooled in preparation for Illumina sequencing with an Illumina Novaseq 6000 by the University of Utah High-Throughput Genomics Shared Resource. All sequencing runs produced 150-bp paired-end reads.

### 4.5 Variant calling

We used *FastQForward* (Carson Holt and Mark Yandell, available upon request) to perform alignment, SNP calling, and filtering of our data to produce genome variant call format (gVCF) files. *FastQForward* is a wrapper for *bwa* (Li & Durbin 2009) and sentieon, which is a faster reimplementation of the polymorphism calling algorithm used by *GATK* (McKenna et al. 2010). *FastQForward* dramatically speeds up the performance of both programs but does not produce output differing from that typical of these programs. Using *FastQForward*, we generated gVCF files for each of the 148 pooled samples and each of the sequenced individuals from the 36-month timepoint. The gVCF files from individuals were then aggregated to build VCF files for each population.

For most analyses, we combined variant data from all the individuals within each 36-month sample to simulate pooled data and allow direct comparisons with the pooled data from other time points. To use the individual sequencing data in these grand analyses, we wrote a custom program that calculated allele counts from the VCFs representing the individual samples, using the genotype at each SNP for each individual to produce allele counts analogous to those available from pooled sequencing in all other populations. The script is available as part of the *VCFStats* package (https://github.com/jgbaldwinbrown/vcfstats). Briefly, for each population, this script counts the two alleles at each locus for each individual, then generates a new VCF file with one column for each population. For example, in a three-individual population with genotypes at a single locus of GA, GG, and GA, the column representing the allele counts in the final VCF file would have a count of 4 Gs and 2 As. These re-formatted data were then combined into an experiment-wide VCF that also included the pooled populations.

For analysis of allele frequencies with *PoolSeq* (Taus et al. 2017) and our custom code, we converted VCF files into synchronized files using the custom script vcf2sync.py, also included in *VCFStats*. This conversion takes allele counts directly from a VCF file, identifies the reference and highest frequency alternative allele for each locus, and generates a sync file containing allele counts for all biallelic SNPs.

### 4.6 Population structure

We used *popvae* (Battey et al. 2020) to identify population structure across the experiment. We applied *popvae* to a VCF file containing SNPs from our dataset with a minor allele frequency *≥* 10% and a coverage greater than 10X in at least one sample. To avoid the problem of correlation between sites due to linkage, we wrote a custom script (*space_vcf* in *vcfstats*) that retains only SNPs spaced greater than a minimum distance (here, 10kb) from each other.

### 4.7 Linkage disequilibrium

We calculated reduction in linkage disequilibrium (LD) *a priori* based on the number of generations from the start of the experiment according to the classical equation:

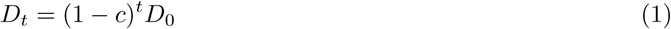

where D is linkage disequilibrium, t is the number of generations, and c is the recombination rate (Slatkin 2008). Because most phenotypic change occurred early in the experiment, we expect the LD subsequent to major allele frequency change to have decayed approximately as many generations as the experiment ran. We further calculated the distance at which decay reduced LD to a given fraction of its initial value as:

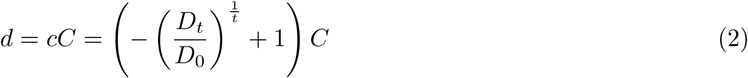

where *d* is the distance producing the given recombination rate and *C* is the chromosome size (assuming 1 recombination event per chromosome, per generation).

### 4.8 Tests for selection

We performed two tests for selection, both by testing for allele frequency differences between populations. We first used the Cochran-Mantel-Haenszel test (Cochran 1954; Mantel & Haenszel 1959) to compare different selection treatments at the 36-month timepoint. We also used the p*F_ST_* test (Kronenberg 2015; Domyan et al. 2016; Garrison et al. 2021) to compare these same treatments, and to compare allele frequencies within a single replicate (https://github.com/vcflib). The parameter p*F_ST_* detects allele frequency differences between populations using a modified likelihood ratio test that incorporates genotype likelihood information. We generated 10-kb sliding window *p*-values from the *CMH* and p*F_ST_* tests using the Fisher method (Fisher 1928). We chose this window size conservatively based on the fact that recombination should yield haplotypes that are, on average, approximately 377 kb long after 48 generations (the 36-month time point is approximately 45 generations in this experiment) (Supplementary Table 1, and see “Linkage Disequilibrium” in Methods). All sliding window analyses were performed with a step size of 50 kb. All-values were corrected by Benjamini-Hochberg FDR (Benjamini & Hochberg 1995). We identified genome-wide effective population size (*N_e_*) using the method “P.planII” in *PoolSeq*’s *estimateNe* function (Taus et al. 2017).

### 4.9 Other statistics

We compared allele frequency slopes between populations of lice in different treatments with Tukey’s Honest Significant Difference test (Tukey 1949). We used Fisher’s Exact test (Fisher 1922) to compare the rate of pairwise overlaps of putatively selected genomic regions for louse populations among the four replicates. For example, the four values in the contingency table in a comparison of two treatments were: a) total significant sites in treatment 1; b) total pairwise overlapped significant sites between any two replicates in treatment 1; c) total significant sites in treatment 2; d) total pairwise overlapped significant sites between any two replicates in treatment 2.

### 4.10 Tests of overlap

We used a permutation approach to identify overlap between genomic regions identified as significant by p*F_ST_*in different replicates of the experiment. We permuted the position, but not size, of each significant region in each dataset, then counted the proportion of permuted sets with more overlap than that of the empirical data. This was our *p* value for overlap. We used 2,000 permutations for each comparison. The custom tool for permuting these intervals, *Permuvals*, is available at https://github.com/jgbaldwinbrown/permuvals.

### 4.11 Allele frequency trajectory calculations

We used the custom tool *tsplot* (https://github.com/jgbaldwinbrown/tsplot) to calculate and plot allele frequency trajectories over the course of the experiment. We used this same tool to calculate lines of best fit and slopes for all trajectories using the formula allele frequency~time. The *tsplot* tool uses the regression package (https://github.com/sajari/regression) for fitting lines of best fit and then does all other calculations, including identifying the selected allele for polarizing plots, collecting alleles from matching sequencing runs for time series generation, identifying the time interval of greatest allele frequency change, and instantaneous slope calculation.

### 4.12 Gene ontology enrichment

We used the default settings of *goatools* (Klopfenstein et al. 2018) to identify enrichment in gene functional categories under peaks of differentiation between populations of lice on birds of different color and preening treatments. Gene Ontology (GO) term associations with genes were drawn directly from the *Columbicola columbae* genome annotation (Baldwin-Brown et al. 2021). All GO terms originate from InterProScan (Jones et al. 2014), and were derived by running InterProScan on the transcripts identified by MAKER (Cantarel et al. 2008).

### 4.13 Functional characterization of polymorphisms

We used *snpdat* (Doran & Creevey 2013) to characterize SNPs as synonymous, nonsynonymous, or untranslated. We used the genome annotation and the filtered polymorphisms described above to identify the set of putative selected, non-synonymous SNPs.

## 5 Results and Discussion

### 5.1 Genomic sequencing and variant calling

To study the genomic basis of adaptive diversification, pigeon lice were experimentally evolved on different color backgrounds by maintaining selection lines on pigeon breeds that differed in color (black, white, and ancestral grey). We investigated allelic variation among the experimentally evolved lice by whole-genome sequencing. We generated a total of 13.8 Tb of whole genome sequencing data. We sequenced 321 pools of lice, to an average of 84.9X genomic coverage, and 2151 individual lice to an average of 18.3X genomic coverage. Of these reads, 11.6 Tb mapped successfully via *bwa*, and polymorphism calling with *FastQForward* identified 4.57 million polymorphic sites based on these reads.

We used *popvae* (Battey et al. 2020) to identify structure in the evolved populations based on individuals sequenced at the 36-month time point. Using a whole-genome assay of population structure, we found that individuals were most closely related to other individuals from the same population; i.e., lice among the 4 pigeons in a single aviary were each other’s closest relatives (Fig. 3). Moreover, populations tended to cluster with the 5 other populations from the same replicate. This result is consistent with the design of the experiment, in which lice transferred to birds in each replicate set of 6 aviaries were drawn from the initial ancestral population 6 weeks after the previous replicate. A common problem with experimental evolution studies is that accidental migration between populations can confound the inference of selection because similarities driven by accidental gene flow produce the same signal of allele frequency similarity as the experimental treatments. In our study, physical isolation of lice between aviaries (see methods), as well as the results from *popvae*, show that accidental migration seldom, if ever, occurred.

**Figure 3:**
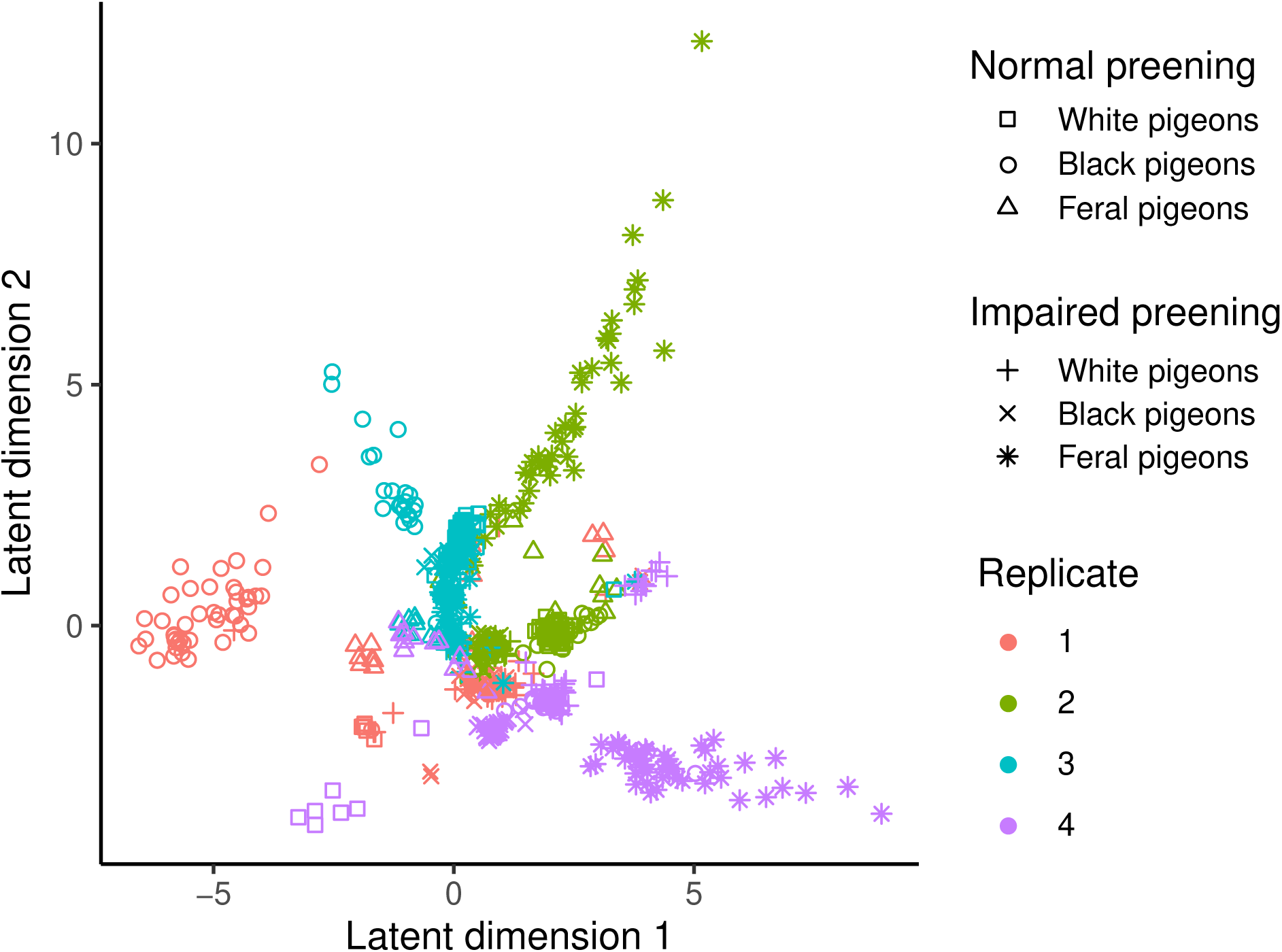
Population structure of experimental populations. Popvae analysis shows that lice within a population – i.e. lice on birds inhabiting the same aviary (same replicate, host color, and preening regime) – are generally more genetically similar to one another, and that populations in the same replicate (Table 1) are more closely related to one another, than they are to lice from the same treatment in other replicates. The lice here were all drawn from populations at the 36-month time point and individually sequenced.

### 5.2 Many genomic regions drive adaptation

A previous study showed that color change in lice on pigeons of different plumage colors is rapid and substantial (Bush et al. 2019). Nevertheless, the genomic signatures of color change in our experimental populations of lice are highly complex and do not include regions that harbor known melanin pathway genes. We searched for loci under selection by comparing allele frequencies between populations using p*F_ST_*(Kronenberg 2015; Domyan et al. 2016; Garrison et al. 2021). We compared allele frequencies at 36 months among lice on normally preening white and black pigeons to the equivalent lice on normally preening grey pigeons (controls). We made this comparison in two ways. First, we took all lice on normally preening white or black pigeons and compared them to all lice on normally preening grey pigeons (i.e., lice from a given treatment were pooled across all four replicates). Second, we compared lice from normally preening white or black pigeons to lice on normally preening grey pigeons within each replicate. In principle, lice on grey (control) pigeons as well as lice on white and black pigeons should be subject to the same “laboratory” conditions, but only lice on white and black pigeons with normal preening should experience selection for cryptic coloration. We controlled our contrasts to only identify selection associated with preening by ignoring all putatively selected loci that were also identified as putatively selected in a matching comparison in preening-impaired birds. In total, we found 103 putative selected sites across our populations, 71 of which came from lice on white pigeons and 32 from lice on black pigeons (Fig. 4). Of the 71 sites from lice on white pigeons, 20 sites were shared with two of the four replicates, while only 3 sites from lice on black pigeons were shared by two replicates (Fig. 4). No sites were shared by 3 or 4 replicates in either the black or white treatments.

**Figure 4:**
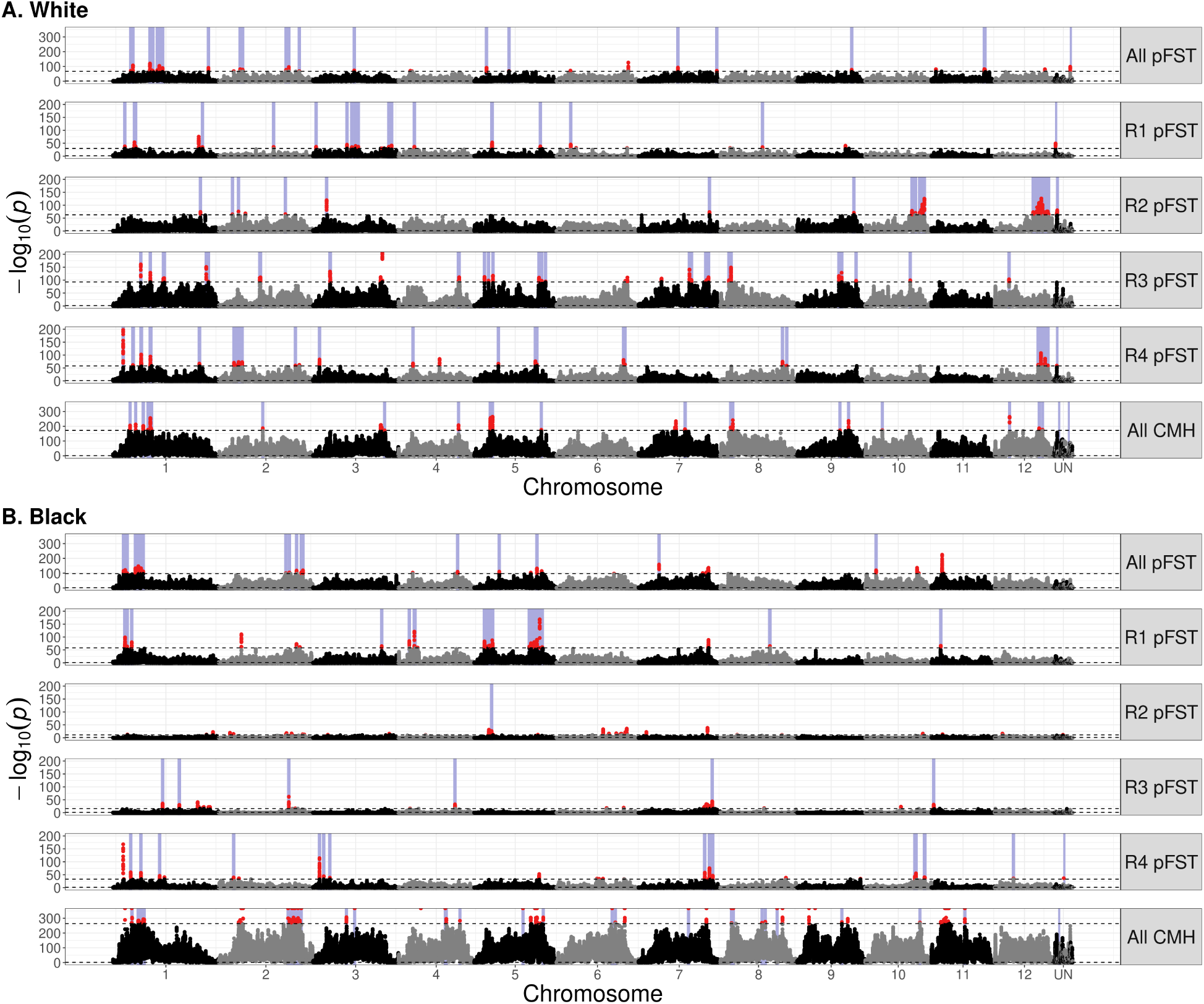
GWAS analyses identified different genomic regions under putative selection in different treatments and replicates. We used p*F_ST_* at month 36 of the experiment to identify regions of the genome under selection. (A-B) We contrasted an experimental population – lice on (A) white pigeons or (B) black pigeons – with lice from grey controls. The top 0.1% of sites (upper dotted line) are shown in red. FDR-corrected −log(p) values are plotted, with the FDR-corrected significance threshold of p = 0.05 near the bottom of each plot (lower dotted line). We calculated p*F_ST_* in each of the four replicates individually (R1-R4), and in a combination of all replicates (all p*F_ST_*). In addition, we used the Cochran-Mantel-Haenszel (CMH) test to identify sites with replicated allele frequency change across replicate populations of the same treatment. Purple bars represent regions where the normally-preening louse population is above the 0.01% statistical threshold, while the impaired-preening louse population on the same color birds is not above 0.01%.

### 5.3 Different loci are selected in different replicates of the same treatment

A major question in evolutionary biology (and experimental evolution studies) is, how repeatable is evolutionary change? That is, if we could turn back the clock and let a population evolve in the same environment a second time, would the same alleles change to adapt to that environment? We tested this idea by comparing the putative selected loci in different replicates of the same selection treatment. This comparison tested whether the same loci were selected repeatedly as an adaptive response to identical selective regimes. Putatively selected sites in many pairs of replicates did overlap between two replicates more than expected by chance (Supplementary Table 2); however, none of the putatively selected loci were shared by three or four replicates (Fig. 5). In fact, most of the putative selected sites are unique to just one replicate in a treatment.

**Figure 5:**
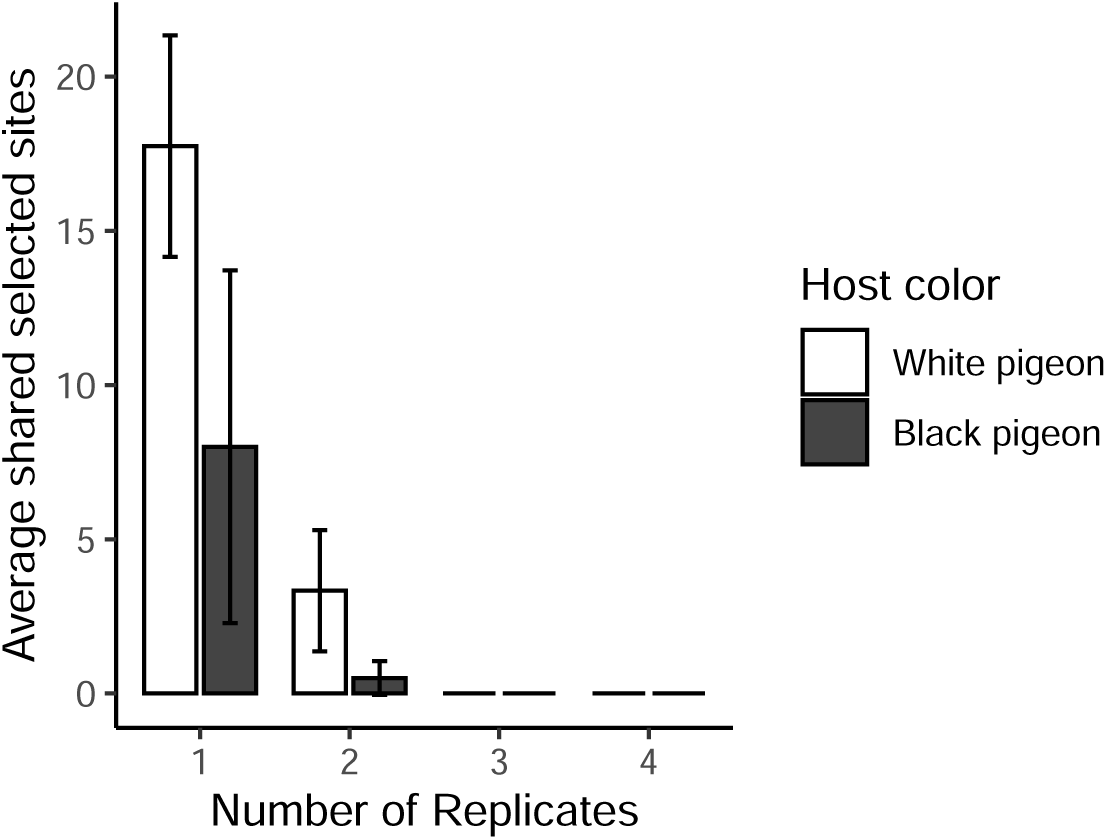
Most selected sites are private to each treatment and replicate. This plot shows the average number of putative selected sites that are private (x-axis = 1) versus shared between replicates (x-axis *≥* 2). While a few putative selected sites were found in two replicates, none were shared by three or four replicates.

A tempting explanation for the overall low overlap between putative selected sites across the different replicates is the balance of selection and genetic drift. Classical population genetic theory argues that selection dominates over drift when selection is strong (*s >* 1*/N_e_*, where *s* is the selection coefficient and *N_e_*is effective population size), but drift dominates when selection is weak (*s <* 1*/N_e_*). If we imagine that all loci contribute to a trait equally, we should expect selection to be detectable when *s/L >* 1*/N_e_*, where *L* is the number of loci contributing to the trait under selection. In this experiment, where sites are largely not replicated, *N_e_* is relatively small, while *L* appears to be large. This is the case in other artificial selection experiments performed on complex traits, such as size, that involve macroscopic plants or animals that are difficult to maintain in large numbers (Mueller et al. 2018, Turner et al. 2011, Turner and Miller 2012, Orozco-Terwengel et al. 2012). Because prior studies have found color to be a relatively complex trait (True 2003; but see other *Drosophila* color mutants such as *ebony* and *yellow*, Wittkopp 2002), we expected to have a high L, and therefore a low rate of overlap, in color evolution. We find no difference between the color selection regimes either in terms of number of selected sites or the chance that selected sites overlap with each other between replicates (Fig. 5, Supplementary Table 3). This result, combined with the large number of loci identified as selected, suggests that the genetic control of color in feather lice is complex. Since most sites that change in allele frequency in response to selection are not shared among replicates, this suggests that chance factors, such as differences in allelic representation in the starting populations for each replicate, also influence the outcome of evolution in our experiment.

Another explanation for the large number of unshared putative selected sites could be a low *s* or a low *N_e_*. Although we cannot directly measure the selection coefficient for color in these experiments, phenotypic evidence from the time course of phenotypic change suggests that selection was strong (Bush et al. 2019). We do, however, have direct measurements of *N_e_* both by counting lice removed from birds by CO2 and by population genetics. These data indicate effective population sizes that range from dozens to hundreds (Supplementary Fig. 1). The small *N_e_* of some of our populations increases the potential role of genetic drift in the evolutionary outcome of the experiment.

### 5.4 Allele frequency trajectories show consistent, ongoing selection

We tracked allele frequencies across the experiment at all polymorphic loci. At each of our putative selected loci, we identified the most significant polymorphic site by p*F_ST_* and plotted its allele frequency over time (Fig. 6). We polarized these sites to measure the frequency of whichever allele is more common in experimental than control populations at 36 months. There is a clear upward trend in allele frequency in these selected alleles over time. This result both supports the inference that these sites are under selection and helps us to characterize the selection that is occurring. While some sites rapidly rise to fixation, most sites change allele frequency gradually over the course of the experiment, with most fixations occurring only toward the end of the experiment (Fig. 7). This is consistent with our phenotypic results, which showed dramatic changes in color early in the experiment followed by continued modest changes later in the experiment. Note that the allele frequencies at these selected sites are biased to be high at 36 months because they were chosen due to their high allele frequency differences between selected and control populations. This causes the frequencies to regress to the mean (Galton 1886) outside of the 36-month time point. We expect allele frequencies at other time points are unbiased.

**Figure 6:**
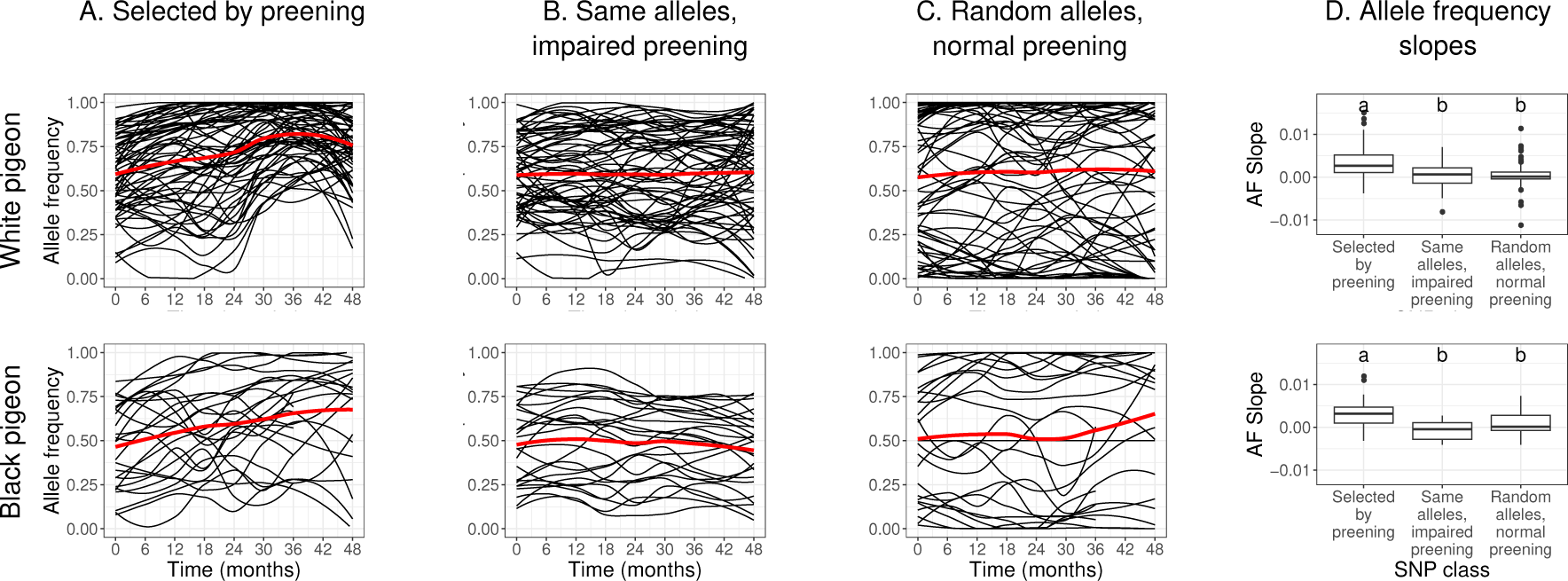
Selected alleles increased in frequency over time. We identified the most significant SNPs by p*F_ST_* in each of our putatively selected loci (65 loci for white, 26 for black), then plotted allele frequencies over time. (A) Allele frequency change in lice from populations exposed to preening-mediated selection (normally preening birds). The red line is the average of all plotted allele frequencies. (B) Allele frequency change at the same sites, but in lice not exposed to preening-mediated selection (impaired preening birds); these populations showed no change in allele frequencies over time. (C) Allele frequency change at randomly chosen sites in louse populations on normally preening birds; these sites have more rare minor alleles than A or B, in keeping with a neutral site frequency spectrum outside of selected regions. (D) Slopes of lines of best fit for the same trajectories in A-C, with the midline representing the median, the box representing the interquartile range, and the whiskers representing 1.5 times the interquartile range. Different lower-case letters indicate statistically significant differences (P<0.01, Tukey’s HSD test).

**Figure 7:**
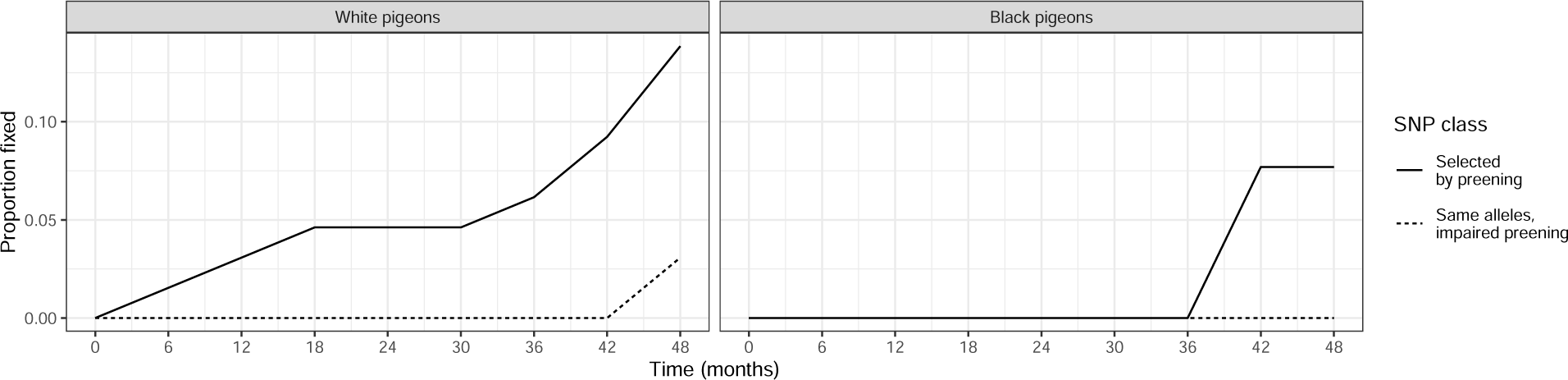
Populations of lice experiencing selection by preening have higher rates of allelic fixation (solid lines) than populations of lice on impaired preening birds (dotted lines). Lines depict cumulative fixations in the polymorphic loci shown in Fig. 6 (65 loci for white pigeons, 26 for black). An allele was assumed to be fixed if its frequency reached 100% and remained at 100% for the remainder of the experiment. Fixations at the putatively selected sites in unselected populations of lice on impaired preening birds were exceedingly rare.

Interestingly, the continued change in allele frequency among louse populations in our study suggests that selection was ongoing and additional adaptation would occur if the experiment were allowed to continue. Further, over macro-evolutionary time, lice in the genus *Columbicola* have adapted to different species of pigeons that vary dramatically in color. For example, the louse species *C. wolffhuegeli* parasitizes the pied-imperial pigeon (*Ducula bicolor*), a species with mostly white plumage. If we extrapolate the rate of phenotypic divergence between white-evolved and grey-evolved lice in our experiment (a difference of about 18 lumens over 4 years), it would take only 14 years to reach the level of phenotypic divergence between *C. columbae* and *C. wolffhuegeli* (65 lumens), which are species of lice that diverged millions of years ago (Johnson et al. 2007, Smith et al. 2011). Though this extrapolation is admittedly simplistic, it implies the potential for very rapid adaptation of feather lice to new hosts.

### 5.5 Differentiated sites in unselected lines do not show evidence of selection

To test for the possibility that the selection we detected was due to factors other than preening-mediated selection, we examined the putatively selected sites in louse populations on pigeons with impaired preening. Specifically, we plotted the frequencies of alleles in these same polymorphic sites from populations of lice on preening-impaired pigeons vs. normally preening pigeons. We also plotted the frequency of alleles in randomly chosen polymorphic sites from populations of lice on normally preening pigeons. The differences between slopes in these trajectories (Fig. 6) are informative. First, the mean slope of selected sites of lice on normally preening populations is significantly higher than the near-zero slope of these same sites in the population of lice on preening-impaired pigeons (Tukey’s HSD test; black p = 5.0×10-6, white *p* = 6.0×10-7), confirming that these sites are indeed under selection in the experimental populations. Second, the mean slope of the putative selected sites is significantly higher than the near-zero slope of randomly-chosen sites from the same population (Tukey’s HSD test; black *p* = 0.0046, white *p* = 2.8×10-6). These results demonstrate that non-random allele frequency change is localized to the regions identified as selected.

### 5.6 Differentiated sites have high starting minor allele frequencies

Site frequency spectra inform us about the types of sites that are likely to respond to selection and likely to be detected as differentiated. The expectation for the site frequency spectrum under neutrality is a large proportion of low-frequency alleles (Kimura 1983). Recent selective sweeps tend to remove polymorphism from the population while raising the allele frequency of the polymorphisms linked to the selected site, thus skewing the site frequency spectrum toward common alleles. The initial site frequency spectra of unselected sites contains a high proportion of low-frequency alleles, as expected in neutral evolution (Kimura 1983) (Supplementary Table 4). In contrast, the selected sites in our analyses have site frequency spectra highly skewed toward common variants in both preening and preening-impaired populations.

Population genetic theory predicts that selection should dominate drift more easily in common variants because they contribute more to the heritability of the trait under selection (Scoville et al. 2011). Thus, common variants are more likely to respond to selection in the first place. Theory also predicts that common variants change allele frequency more rapidly (Zhao et al. 2013), suggesting that power to detect allele frequency change is highest in common variants. Overall, these theoretical expectations present two possibilities for why the allele frequencies at selected sites are skewed toward common alleles. It may be that common alleles are overrepresented at our selected sites because selection removed rare alleles, or it may be that common alleles are overrepresented because common alleles are more likely to respond strongly to selection in the first place. These two explanations are not mutually exclusive and are difficult to distinguish here.

### 5.7 Selection is removing trait-affecting variation from the population

A central question in evolutionary biology is whether adaptation removes useful variation from the population. That is, if a population is selected to change some trait, does it lose the variation needed to change that trait again in the future? To better understand this possibility, we characterized the loss of variance at selected sites. We scored variants as fixed or lost in a population if they reached and remained at an allele frequency of 100% or 0%, respectively (Fig. 7). Allele fixation and loss can be caused by either genetic drift or selection; we found evidence for both. Our putative selected sites were more likely to fix in normally preening populations than in preening-impaired populations. This difference in fixation probability reinforces the hypothesis that the genetic variants associated with the selected phenotype(s) were responding to preening-driven selection. Past experimental evolution studies in insects show that, when selection toward a phenotype is reversed, adaptation in the opposite direction occurs as quickly as initial adaptation, implying that the genetic variation needed to adapt was not lost in these populations, even in experimental evolution that ran for hundreds of generations (Rose 2004). In contrast, we found that approximately 10% of putative selected sites were fixed or lost by the end of our experiment. If this experiment continued, we would expect much of the original variation driving these populations to be lost, making reverse adaptation at these sites much slower than initial adaptation due to a loss of standing variation.

### 5.8 Many more undetected sites are likely under selection

Our method for identifying selected loci focused on just one time point and identified only sites that were significantly differentiated from a control population. An alternative approach is to test for outlier allele frequency trajectories within each population. Selected alleles are expected to change frequency much more quickly than neutral alleles, so allele frequency trajectories with sharp slopes are likely to be the selected ones (cf. Li and Barton 2023). This approach (Supplementary Fig. 2, 3) yields a very different set of selected sites. Although the evidence for selection in these sites is strong (the top 0.1% of 10-kb sliding window averaged allele frequency slopes), there are several reasons to be more skeptical that these sites are under selection.

Unlike p*F_ST_*, which takes sequencing coverage into account and assigns lower confidence to allele frequency differences at sites with low coverage, our slope calculation does not correct for sequencing coverage. Because of this, many slope-outliers are at chromosome edges, where sequencing coverage is poor and allele frequency estimates are less accurate. Also, unlike with p*F_ST_*, it is hard to differentiate a slope-outlier due to experimental preening conditions from a slope-outlier that is due to selection for a trait other than the one of interest. This limitation exists because we did not contrast the slope of alleles in preening populations to slopes in non-preening populations. In summary, while some of the slope-outliers are likely examples of real selection, it is hard to distinguish these from false positives.

### 5.9 Genes in differentiated regions

To better understand the genes and cellular functions that may be involved in selection for color, and to see if adaptation for color in lice resembles color adaptation in other taxa, we characterized the genes in the regions under selection using the gene ontology (GO) tool *goatools* (Supplementary Data Table 1) (Klopfenstein et al. 2018). We found 12 significantly enriched GO terms in our putatively selected sites (Supplementary Table 5). Selected loci in dark-adapted lice were enriched for two cation transporter activity GO terms (cation-transporting ATPase complex, sodium:potassium-exchanging ATPase complex). Because such transporters are involved in many cell functions, they could plausibly be mechanistically connected to color, especially relating to pigment transport and deposition in the body. Melanin is known to have a high affinity for metal cations (Potts and Au 1976, Larsson and Tjälve 1978) and cation transport is implicated in the development and function of melanosomes in vertebrates (Bellono and Oancea 2014, Wiriyasermkul et al. 2020). Still, the connection between cation transport and pigment variation in our louse populations is speculative.

While white-adapted lice showed enrichment for several GO categories, most were connected to nervous system function and olfaction. Our selection regimes did not specifically target olfaction traits, but melanin and the neurotransmitter components associated with olfaction use some of the same precursor molecules (Gelis et al. 2016, Pavan and Dalpiaz 2017). One might imagine these neurotransmitter precursors are under selection in our populations not because of their role in olfaction, but because they also affect color. However, we see no evidence for genes directly related to these molecules in our enrichment set. There are connections between nervous system function and pigment deposition, notably the role of pigment dispersing factor in the regulation of circadian rhythm (Park & Hall 1998; Matsushima et al. 2004). The product of one gene in a significantly differentiated genomic region in lice adapted to white birds, *Pdx*, was shown to have protein-protein interactions with pigment dispersing factor in the *Drosophila* protein interaction map (DpiM; Guruharsha et al 2011).

## 6 Conclusion

Adaptation of parasites to novel hosts, either due to a host switching event or evolution of a host population, is an opportunity for adaptive radiation. We sequenced the genomes of pigeon lice subjected to artificial host switches to characterize the genetic underpinnings of an experimental adaptive radiation. Our results indicate that one trait under selection, louse body color, appears to be genetically complex with numerous contributing loci. We also found that the same loci were rarely involved in replicate lineages of lice exposed to very similar conditions. We used allele frequency trajectories at selected sites to show that adaptation is rapid and selection persisted for the duration of the 48-month experiment (ca. 60 louse generations). We also found that selection causes the loss of some genetic variation, which may make phenotypic reversion more difficult and may reduce the fitness of these lice if they were to encounter grey hosts in the future.

## 8 Animal Ethics Statement

All procedures involving vertebrates were approved by the University of Utah Institutional Animal Care and Use Committee, protocol #14-06010.

## 9 Acknowledgements

We thank André Watson for his help maintaining the louse populations, collecting lice, and physical measurement of lice; and Anna Vickrey for assistance in optimizing DNA extractions.

This work was supported by the National Science Foundation (DEB-0107947 to DHC; DEB-1342600 to DHC, SEB and MDS; DEB-1926738 to DHC and SEB; and DEB-1342604, DEB-1926919, DEB-1925487, DEB-2328118 to KPJ); and the National Institutes of Health (R35GM131787 to MDS). We also thank Nitin Phadnis for his advice and acknowledge the National Institutes of Health grant R01GM141422. The support and resources from the Center for High Performance Computing at the University of Utah are gratefully acknowledged. The computational resources used were partially funded by the National Institutes of Health Shared Instrumentation Grant 1S10OD021644-01A1.

## 11 Supplementary materials

### 11.1 The evolved populations are largely structured by ancestral relationships

We used popvae (Battey, Coffing, and Kern 2020) to identify structure in the evolved populations based on individuals sequenced in the ancestral generation and the 36-month time point. Based on a whole-genome assay of population structure, we found (Fig. 3) individuals could clearly be grouped by their respective populations. Because we ran popvae on a broad set of polymorphisms drawn from across the genome, we expect differentiation of the populations to be driven primarily by genetic drift and population structure (here, founder effects). While selection should act at a subset of loci, drift acts on the entire genome; thus, measures of population structure should be driven more by drift than by selection, excepting cases where selection is occurring across the entire genome.

Notably, the populations cluster according to their replicate number. This is consistent with the design of the experiment. Each replicate set of treatments was drawn from the initial ancestral population in a manner staggered in time. Replicate 2 was founded six weeks after replicate 1, replicate 3 was founded six weeks after that, and replicate 4 six weeks after that. This was done to make handling the lice more practical. Owing to this, lice from a given replicate should be more closely related to other lice from the same replicate, even if they are from another treatment, than they are to other lice within their own treatment, but from a different replicate. This is what is shown in the popvae data.

### 11.2 Supplementary tables

**Supplementary Table 1:** Linkage disequilibrium decays to produce approximately 377kb haplotypes by 48 generations. The expected distance between recombination is calculated by assuming one recombination event per chromosome per generation, then calculating the distance in basepairs expected by a given number of generations, i.e., (genome size / number of chromosomes) / generations. The recombination rate required is the rate of recombination between two points needed to reduce linkage disequilibrium to 1/3 of its initial value. It is equal to 1 *−* (*final*_*ld/initial*_*ld*)^(1^*^/generations^*^)^. The basepair distance corresponding to recombination rate is simply the recombination rate times the chromosome size, and indicates the average distance between two loci that would have a recombination rate such that linkage disequilibrium would be reduced to 1/3 of its original value.

| Generation | Expected distance between recombination events | Recombination rate required | Basepair distance corresponding to recombination rate |
| --- | --- | --- | --- |
| 6 | 2777778 | 0.1673 | 2788845 |
| 12 | 1388889 | 0.0875 | 1458214 |
| 18 | 925926 | 0.0592 | 986900 |
| 24 | 694444 | 0.0447 | 745793 |
| 30 | 555555 | 0.0360 | 599353 |
| 36 | 462962 | 0.0301 | 500979 |
| 42 | 396825 | 0.0258 | 430343 |
| 48 | 347222 | 0.0226 | 377164 |
| 54 | 308642 | 0.0201 | 335682 |

### 11.3 Supplementary figures

**Supplementary Figure 1:**
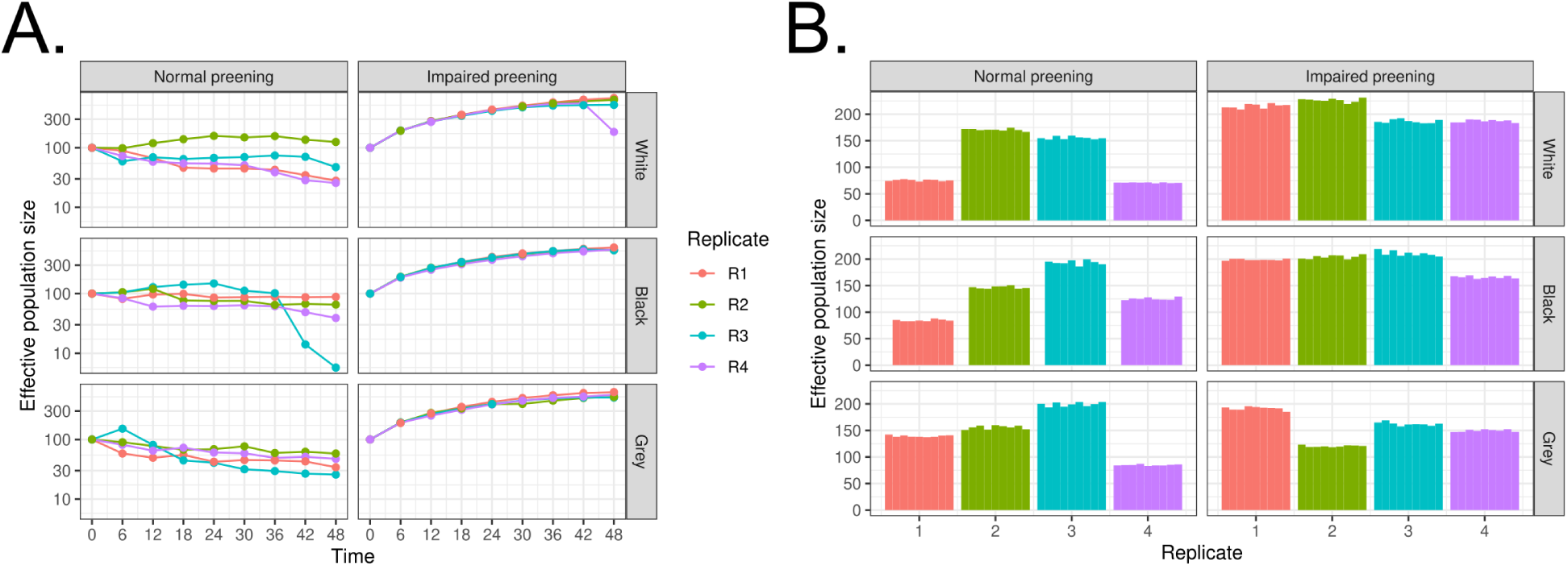
Heterogeneity in effective population sizes across treatments is an important parameter for detecting selection. We estimated effective population size in two ways: (A) The number of lice recovered by CO2 fumigation in each population over the course of the experiment, corrected by harmonic mean to reflect the effective population size. (B) Effective population size, as inferred from the increase in variance of allele frequencies over time.

**Supplementary Figure 2:**
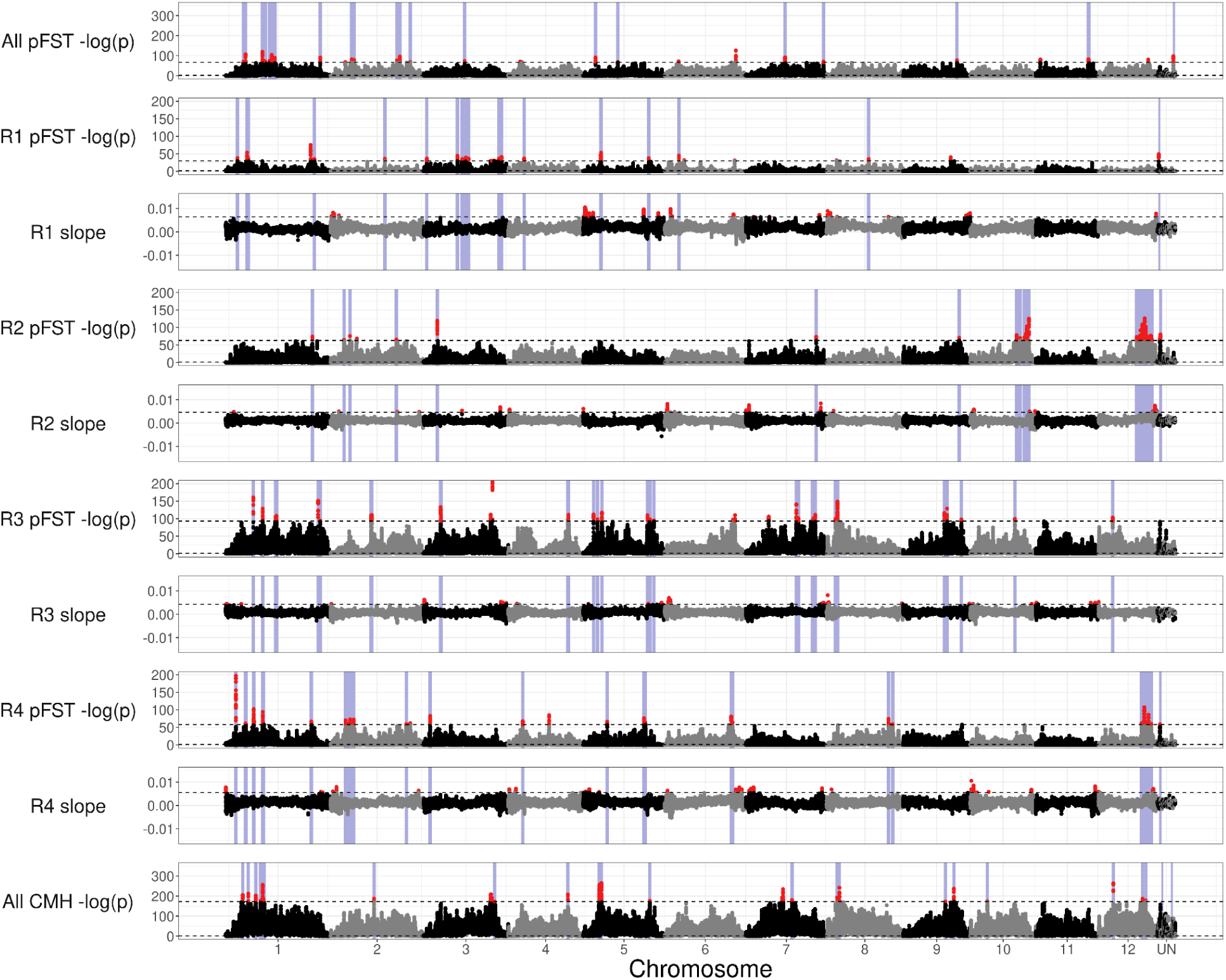
Outlier allele frequency slopes largely do not match p*F_ST_* outliers, showing that allele frequency slopes may be an effective tool for identifying selected regions missed by traditional tests of differentiation. This plot covers populations reared on white pigeons. This plot matches figure 4, but includes the 10kb-windowed allele frequency slope as well. With the caveat that slope calculations do not correct for low-coverage region, many of the outlier regions in the “slope” plots above differ from their matching “p*F_ST_*” plots.

**Supplementary Figure 3:**
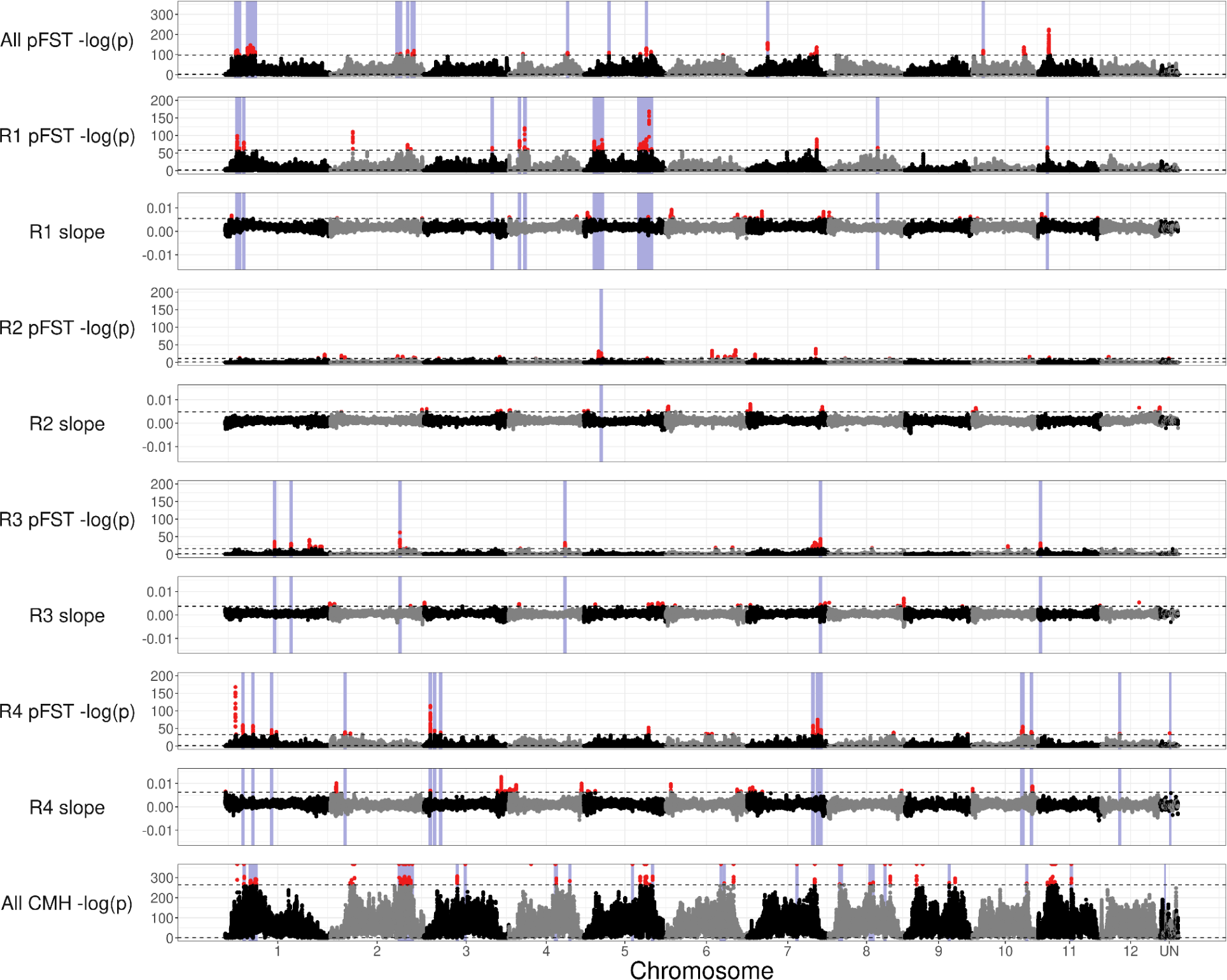
Outlier allele frequency slopes largely do not match p*F_ST_* outliers, showing that allele frequency slopes may be an effective tool for identifying selected regions missed by traditional tests of differentiation. This plot covers populations reared on black pigeons. This plot matches figure 4, but includes the 10kb-windowed allele frequency slope as well. With the caveat that slope calculations do not correct for low-coverage region, many of the outlier regions in the “slope” plots above differ from their matching “p*F_ST_*” plots.

